# Arousal elevation drives the development of oscillatory vocal output

**DOI:** 10.1101/2021.12.13.472498

**Authors:** Yisi S. Zhang, John L. Alvarez, Asif A. Ghazanfar

## Abstract

Adult behaviors, such as vocal production, often exhibit temporal regularity. In contrast, their immature forms are more irregular. We ask whether the coupling of motor behaviors with arousal changes give rise to temporal regularity. Do they drive the transition from variable to regular motor output over the course of development? We used marmoset monkey vocal production to explore this putative influence of arousal on the nonlinear changes in their developing vocal output patterns. Based on a detailed analysis of vocal and arousal dynamics in marmosets, we put forth a general model incorporating arousal and auditory-feedback loops for spontaneous vocal production. Using this model, we show that a stable oscillation can emerge as the baseline arousal increases, predicting the transition from stochastic to periodic oscillations observed during marmoset vocal development. We further provide a solution for how this model can explain vocal development as the joint consequence of energetic growth and social feedback. Together, we put forth a plausible mechanism for the development of arousal-mediated adaptive behavior.

## Introduction

Over the course of development, the transformation from variable exploratory motor outputs into structured ones is ubiquitous across different behaviors. For many altricial animals, vocal production is crucial from the first day of life as it is the only means to solicit attention from caregivers [1, 2]. Early vocalizations are driven by internal needs and act as scaffolding for later communicative functions [1]. In the infants of humans [3, 4], songbirds [5, 6], bats [7], and marmoset monkeys [8-10], vocalizations transition through different states on their way to becoming adult-like. While it is natural to presume that a behavioral state change must be the result of a neural change, the fact of the matter is that the development of vocal behavior involves multiple components, including the biomechanics of the vocal apparatus, the metabolic energy, neural systems, and the social environment [11]. These moving parts interact, form multiple feedback loops, and self-organize into identifiable structures and functions.

Vocalizations, not only during infancy but also through adulthood, are used to coordinate activities among individuals that are crucial for survival. They are tightly correlated with internal states such as arousal [12-16]. Physiological arousal is associated with global changes in brain and body states that modulate sensory processing and allocate metabolic energy to prepare the body for actions [17]. Arousal levels are correlated with a number of acoustic features in marmoset vocalizations, with higher arousal associated with longer, louder, and less noisy sounds [18]. Arousal is also related to the sequential organization of vocalizations on a longer timescale [8, 12, 18]. Spontaneous vocal sequences of infant and adult marmosets exhibit a high coherence with heart rate fluctuations around 0.1 Hz. In songbirds, males in a high arousal state induced by the presence of females produce more stereotyped songs than in the undirected context where the songs are more variable [19]. How arousal interacts with the vocal production system to affect the regularity of vocal output is unknown.

In this study, we link these two phenomena of 1) the transitions from stochastic to structured behaviors during development and 2) the arousal correlated vocal sequence organization to understand the pattern of shifts in marmoset vocal development. A useful approach for investigating transitions between behavioral states occurring across development is the application of dynamical systems theory [20]. Stable motor dynamics at timescales of seconds can be understood as consequences of control parameters that change over days and months [21]. The control parameters are biological factors that drive the vocal output into new modes as they cross critical points to enter new states [11, 22-24]. Based on behavioral and physiological data, we describe here a parsimonious model incorporating the vocal production system with arousal dynamics. Using this model, we predicted nonlinear changes in vocal sequences during the development of marmosets and tested these predictions with empirical data. We then discuss the model’s insights and offer interpretations of the control parameters.

## Results

### Vocal output and arousal fluctuation are dynamically coupled

Infant marmosets start vocalizing from the first postnatal day. In a context in which the infants were briefly isolated, they vocalized spontaneously and continuously (data published previously [10, 25]). The initial vocalizations were babbling-like, composed of a variety of acoustic patterns, including both mature and immature-sounding calls (e.g., postnatal day 3 (P3) vocalizations; Fig. 1A). Over the first two months, vocalizations became sparser and more regular, dominated by a tonal, long-duration contact call (example vocalizations highlighted by red boxes in Fig. 1A), which is also the vocalization used for long-distance communication in adults.

**Figure 1.**
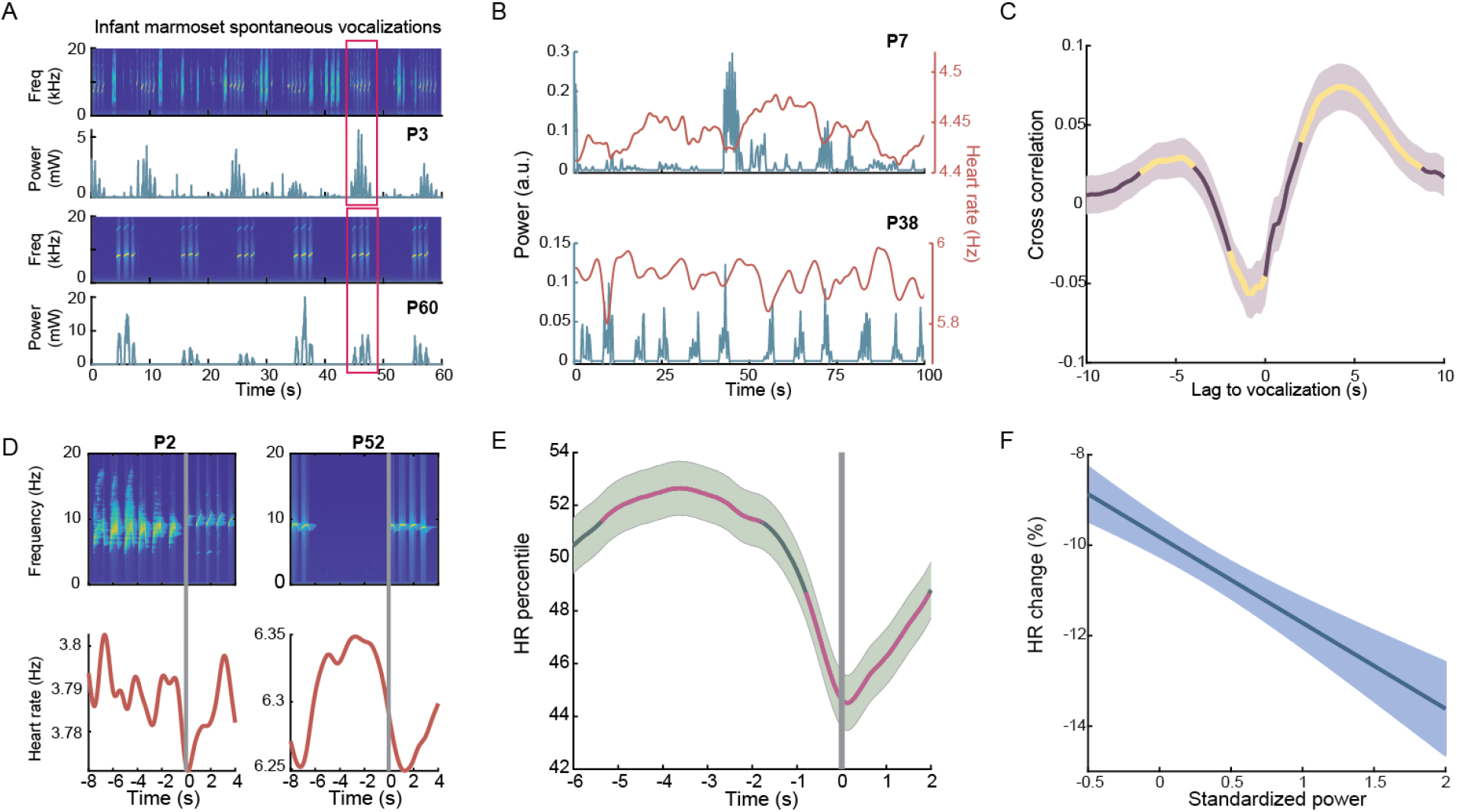
Temporal coordination between vocal output and arousal fluctuations. (A) Two examples of infant vocalizations on two postnatal days. The top two panels are the spectrogram and power of sound amplitude envelope on P3, and the bottom two are on P60. Red boxes highlight contact calls. (B) Two examples of infant vocalization amplitude envelope with real-time heart rate. (C) Cross-correlation between sound and heart rate. The highlighted segments are significantly different from 0 (p<0.05). Shaded area is 95% CI of data. (D) Two examples of heart rate change around the onset of tonal contact calls. (E) Heart rate change around contact call onset at the population level. Heart rate data were converted to percentiles of the session. Highlighted segments in magenta are outside 95% CI of shuffled data. (F) Linear fitting of largest heart rate drop from call onset vs. acoustic power of the call standardized per session.

To quantify the temporal structure of vocal output over hundreds of seconds, we calculated the sound amplitude envelope that smoothed out the fine structure of the acoustic signals (Fig. 1B). Previous studies identified high coherence around 0.1 Hz between spontaneous vocal output and momentary arousal levels (using heart rate as a proxy; see Fig. 1B) in both infants and adults [8, 18], suggesting a robust relationship between these two signals regardless of age. This timescale is much slower than the respiratory rhythm (which is at ~1 Hz), and thus the structural changes at low frequencies are likely to be attributed to arousal but not respiratory development. We next investigated the temporal coordination between vocalization and arousal in detail.

Cross correlation analysis between vocalization and heart rate exhibited a significant dip around 0 and two peaks about 10 s apart, indicating an out-of-phase relationship with an approximately 10-second period (n=7 subjects, P1-P61; t-test with Bonferroni correction; Fig. 1C). To visualize arousal change around vocal production, we aligned the heart rate signals with the onsets of contact calls (Fig. 1D). We observed a general trend that heart rate was significantly high (>50 percentile of session heart rate) before call onsets and dropped significantly upon call production (Fig. 1E; n=2972 contact calls, P1-P61; shuffle test). Considering all types of vocalizations, we found that louder sounds led to a greater reduction of heart rate (Fig. 1F; n=6763 calls, p=4.5e-9, multiple linear regression (MLR)). These findings have three implications: 1) arousal is discharged upon vocalization (Fig. 1F); 2) arousal automatically builds up (Fig. 1E); 3) arousal levels gate vocal production (Fig. 1E). Based on these observations, we offer a model to describe these processes.

### A dynamic model for spontaneous vocal production

Infants often produce exploratory movements that increase in structure over time [26]. What refines those movements into structured behaviors is often reinforcement signals in the form of sensory feedback [27-30]. This is true for the specific case of vocal development, as well [31, 32]. The positive sensorimotor feedback loop is crucial for sustained motor activities [33]. In our model, we assumed that the loudness of self-generated vocalization is reinforced and provides delayed feedback to drive the vocal system (Fig. 2A). On the other hand, positive feedback loops are intrinsically unstable. An antagonistic process is that once the action (vocal production) is fulfilled, arousal is discharged – similar to the idea of Lorenz’s psycho-hydraulic model [34](Fig. 2A). Thus, three dynamical processes constitute our phenomenological model: 1) the neural activity of the motor area *v*(*t*) that generates vocalizations; 2) the vocal signal *s*(*t*) providing positive self-generated auditory feedback; and 3) the arousal level *a*(*t*) that modulates the gain of the drive and is discharged by vocal production.

**Figure 2.**
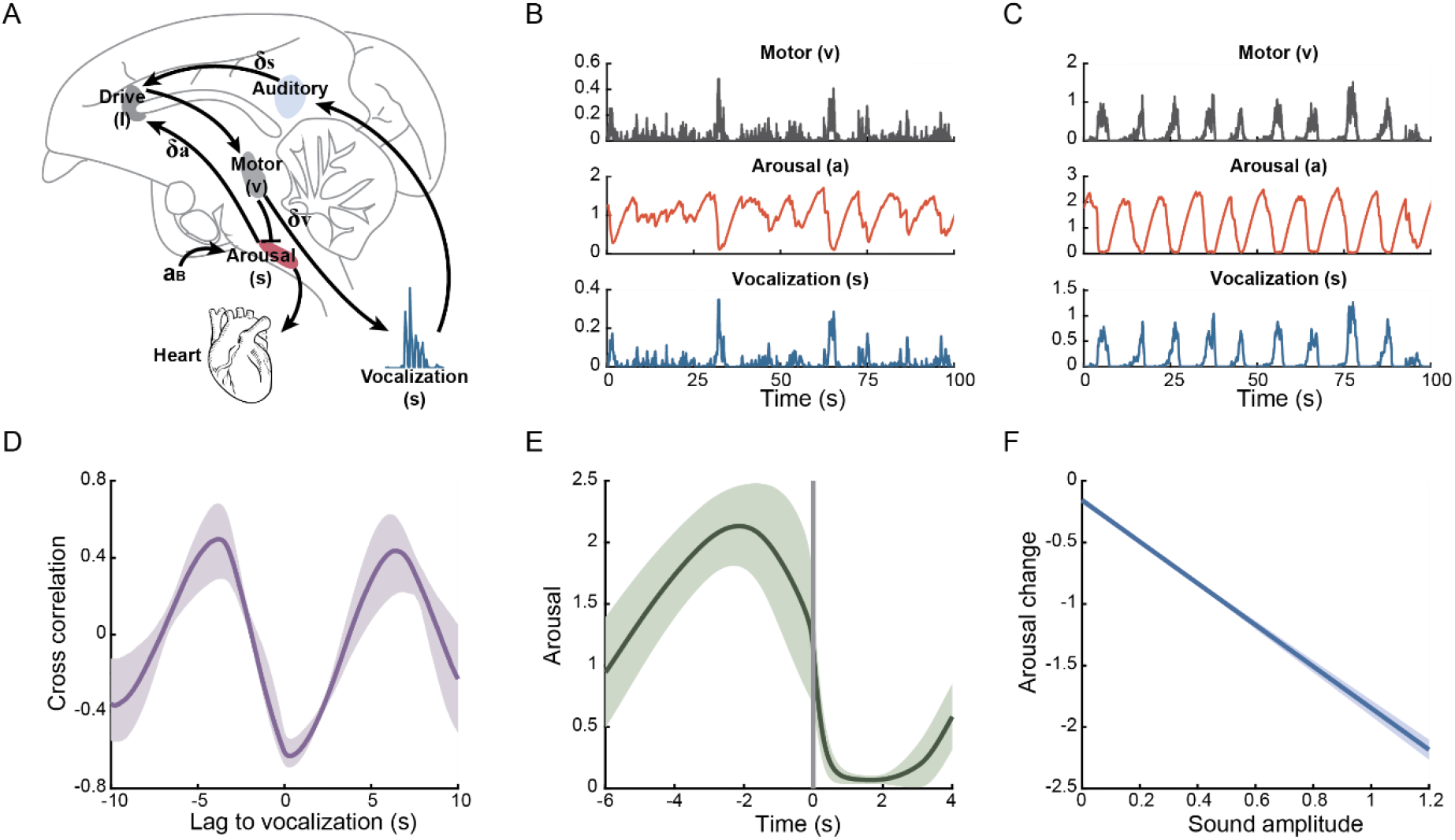
Model setup and simulation. (A) Diagram of model setup. The drive (*I*) integrates delayed arousal (*a*) with excitatory input from auto-auditory feedback (delayed *s*). The motor area (motor, *v*) receives input from the drive and produces vocalization (*s*). The motor signal inhibits arousal, decreasing heart rate. The arousal can be restored to a baseline level *a*_*B*_. *δ*_*a*_, *δ*_*s*_, and *δ*_*v*_ are the latencies of the signals. The anatomical locations are illustrative and not exact. (B) Simulated data for each compartment of the model. From top to bottom are simulations for *v*(*t*), *a*(*t*), and *s*(*t*). This is a simulated example of early vocal output. *a*_*B*_ = 3.1, and the other variables are set as in Table 1. (C) A simulation of later vocal output. *a*_*B*_ = 5.5. (D) Cross correlation between vocalization *s*(*t*) and arousal *a*(*t*). (E) Arousal dynamics around the call onsets (selected calls are >2 s duration). (F) Linear fit between mean sound amplitude of a call and the largest arousal reduction from call onset within a call. For (D-F) *a*_*B*_ ∈ [3, 6]. Shaded areas are the 95% CI of the estimate.

Motor activity generates the neural signal for vocal production. Biologically, this may correspond to an area (e.g., the periaqueductal gray (PAG)), upstream to the brainstem pattern generators. The motor activity *v*, describing a population level firing rate, is driven by a drive signal *I*. *I* is modified by an s-shaped activation function *f* as an input to *v* (Eq. 1). On top of the deterministic dynamics of *v*, there is a stochastic component simulating the intrinsic fluctuation of the neural population, which includes Gaussian noise with an activity-dependent amplitude 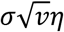. The drive *I* integrates self-generated auditory feedback (the animal’s own vocalization) with the momentary arousal state. This process may take place in the anterior cingulate cortex (ACC) where strong reaction to socially relevant auditory input and vocal production are found [35] and is involved in cardiovascular regulation [36, 37]. We modeled *I* as delayed auditory feedback *s*(*t* − *δ*_*s*_) plus a constant bias *h*, multiplied by the gain of a delayed arousal level *a*(*t* − *δ*_*a*_) (Eq. 2). The vocal output *s*(*t*) is modeled as low-pass filtered muscle activation, which is proportional to the neural activity with a time delay *δ*_*v*_ [38]. The arousal level *a*(*t*) is depleted by the vocal action and slowly replenished to a baseline level *a*_*B*_. The rate of arousal reduction is proportional to the strength of the neural input, compatible with the observation that louder sounds reduce arousal more than quieter ones (Fig. 1F). The arousal state is regulated by several subcortical areas that are responsible for maintaining internal physiology and also modulating cortical activity. We modeled the ascending influence on the drive, e.g., via a pathway to ACC, as a delayed arousal signal *a*(*t* − *δ*_*a*_). The interplay among neural activity, vocal output, and arousal levels together determines the vocal pattern. The complete schematic of this system is shown in Fig. 2A, with equations listed as

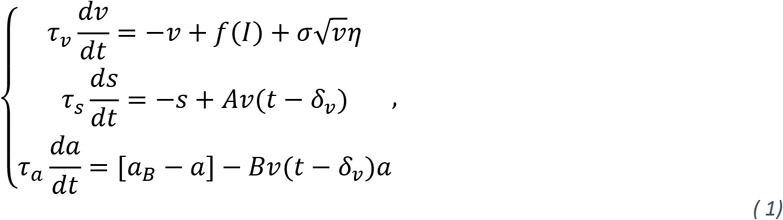

where

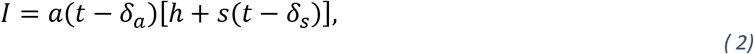

*f* is an s-shaped function taking the form 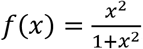, *v*, *s*, *a* are the time-dependent variables of neural activity, sound, and arousal level respectively, τ_*v*_, τ_*s*_, τ_*a*_ are the corresponding time constants, *A* and *B* are positive parameters, η(*t*) is Gaussian noise (zero mean unit variance) scaled by an activity-dependent amplitude 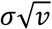. Parameters are given in Table 1.

**Table 1.** Parameters

| $A$ | $B$ | $\tau_v$ | $\tau_s$ | $\tau_a$ | $\delta_v$ | $\delta_s$ | $\delta_a$ | $a_B$ | $\sigma$ | $h$ |
| --- | --- | --- | --- | --- | --- | --- | --- | --- | --- | --- |
| 1 | 130 | 0.1 | 0.1 | 8 | 0.01 | 0.02 | 3 | 0-10 | 1 | 0.1 |

We show that this model can generate signals that resemble the amplitude envelope of marmoset vocal output. By varying the parameters, we simulated the temporal patterns corresponding to early (Fig. 2B) and later (Fig. 2C) developmental stages, comparable to the P3 and P60 examples in Fig. 1A. Essentially, a higher degree of temporal regularity appears later in life. The simulated data show that the cross correlation between sound and arousal signals is qualitatively similar to that observed in Fig. 1C due to the out-of-phase relationship (Fig. 2D). The model also captures the arousal dynamics, which increase before vocal onsets, dip upon vocal production, and recover afterward (Fig. 2E). Furthermore, the model captures the correlation between arousal reduction upon vocalization and the corresponding sound amplitude (Fig. 2F). Thus, this model can produce similar temporal coordination between vocalization and arousal fluctuation as the empirical data.

### Changes in a parameter correlate with shifts in vocal dynamics

Parameter changes can shift the behavior of the system. Among the parameters, we assume that the time-related ones, namely, τ_*v*_, τ_*s*_, τ_*a*_, *δ*_*v*_, *δ*_*s*_, and *δ*_*a*_, are determined by intrinsic neurophysiological properties and remain relatively constant throughout development. In fact, the median frequency of sound amplitude envelope did not change significantly across postnatal days (Fig. 3A; p=0.50), and the frequency is primarily determined by these parameters based on the model (Appendix B). We also assume that the parameters *A* (the rate of neural signal translated into vocal output), *B* (the rate of arousal reduction upon vocal production), and the bias *h*, are unchanged. We, therefore, focus on the role played by the baseline arousal *a*_*B*_. To see how *a*_*B*_ may change during development, we fitted *s* and *a* in our model to the real vocalization and heart rate data. We used an unscented Kalman filter for simultaneous state and parameter estimation of *v*(*t*), *s*(*t*), *a*(*t*), and *a*_*B*_ while fixing other parameters [39] (Fig. S1). Because the state variables *s* and *a* are unit-less, we normalized the sound amplitude and scaled the heart rate for each session with an optimized scaling factor such that the tracking error of vocalization and heart rate was minimized (Methods). The data fitting shows that the estimated *a*_*B*_ increased significantly with postnatal days (p=4.7e-10, MLR; Fig. 3B).

**Figure 3.**
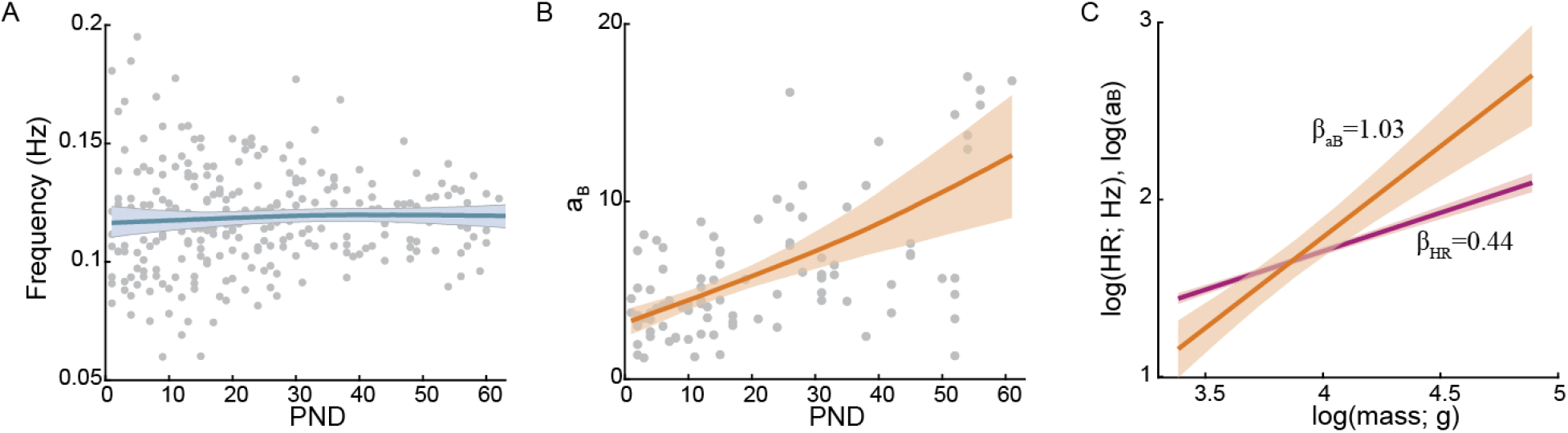
Parameter estimation for *a*_*B*_. (A) Median frequencies of the sound amplitude envelope. Each point is a session. Blue line is the cubic spline fitting of the frequency. (B) Fitted *a*_*B*_ using unscented Kalman filter as a function of postnatal days (PND). Each dot is a session. Red line is cubic spline fitting of the estimated *a*_*B*_. Shaded area is 95% CI of the curve fitting. (C) Linear fitting in the logarithmic scale to estimate allometric scaling of *a*_*B*_ and heart rate. *β*_*aB*_ and *β*_*HR*_ are slopes. Shaded areas are 95% CI.

Although we scaled the data for each session, it is possible that the scaling factor was constant, and changes in *a*_*B*_ simply reflect an elevation of baseline heart rate during development. To test this, we estimated the allometric scaling of *a*_*B*_ with body mass and compared it to the scaling exponent of heart rate. The allometric scaling is the power-law relationship between a biological parameter *y* with body mass *m*: *y* = *αm*^*β*^. Infants tripled their body masses during the first two months [10]. If *a*_*B*_ is proportional to mean heart rate, we should expect identical scaling exponents. The scaling exponent for heart rate was *β*_*HR*_ = 0.44 ± 0.02 (*R*^2^ = 0.78, p=1.6e-31), whereas the scaling exponent for *a*_*B*_ was *β*_*aB*_ = 1.02 ± 0.13 (*R*^2^ = 0.41, p=2.0e-11), significantly greater than *β*_*HR*_ (p=1.6e-4; Fig. 3C). Thus, *a*_*B*_ cannot be simply interpreted by baseline heart rate. We will discuss two potential factors that may contribute to the increase of *a*_*B*_ in the last section.

### The model predicts a Hopf bifurcation in vocal development

One prediction of this model is that increasing *a*_*B*_ can drive the system to produce periodic oscillations via a Hopf bifurcation (Appendix A). A Hopf bifurcation can be defined by the behavior of eigenvalues (λ), which are growth rates of trajectories around a fixed point of the system. λ with a positive real part indicates that the trajectory will leave the fixed point while a negative one indicates convergence to the fixed point with time. The bifurcation occurs when a complex conjugate pair of λ becomes purely imaginary. Oscillations arise as the real part turns positive with a non-zero imaginary part and vice versa. As shown in Figure 4A, the three eigenvalues of this 3-dimensional system have negative real parts when 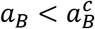, indicating that the system is attracted to a stable fixed point. As *a*_*B*_ crosses 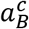, the real part of a pair of complex conjugates becomes positive, enabling oscillations around a frequency of *ω* = *Im* λ. Another critical point lies at a much larger *a*_*B*_, but this is not physiologically plausible (Appendix A, Fig. S2). It can be proven that the oscillation period is on the order of twice as *δ*_*a*_ (Appendix B). We thus set the *δ*_*a*_ such that oscillations of ~0.1 Hz are produced (Fig. 4B). Figure 4B also shows that the frequency deviates minimally with *a*_*B*_ from the birth of oscillation. This is consistent with our observation in Figure 3A that the frequencies were not significantly changed with postnatal days. Such a feature supports a Hopf bifurcation where oscillations are born with finite frequency and zero amplitude.

**Figure 4.**
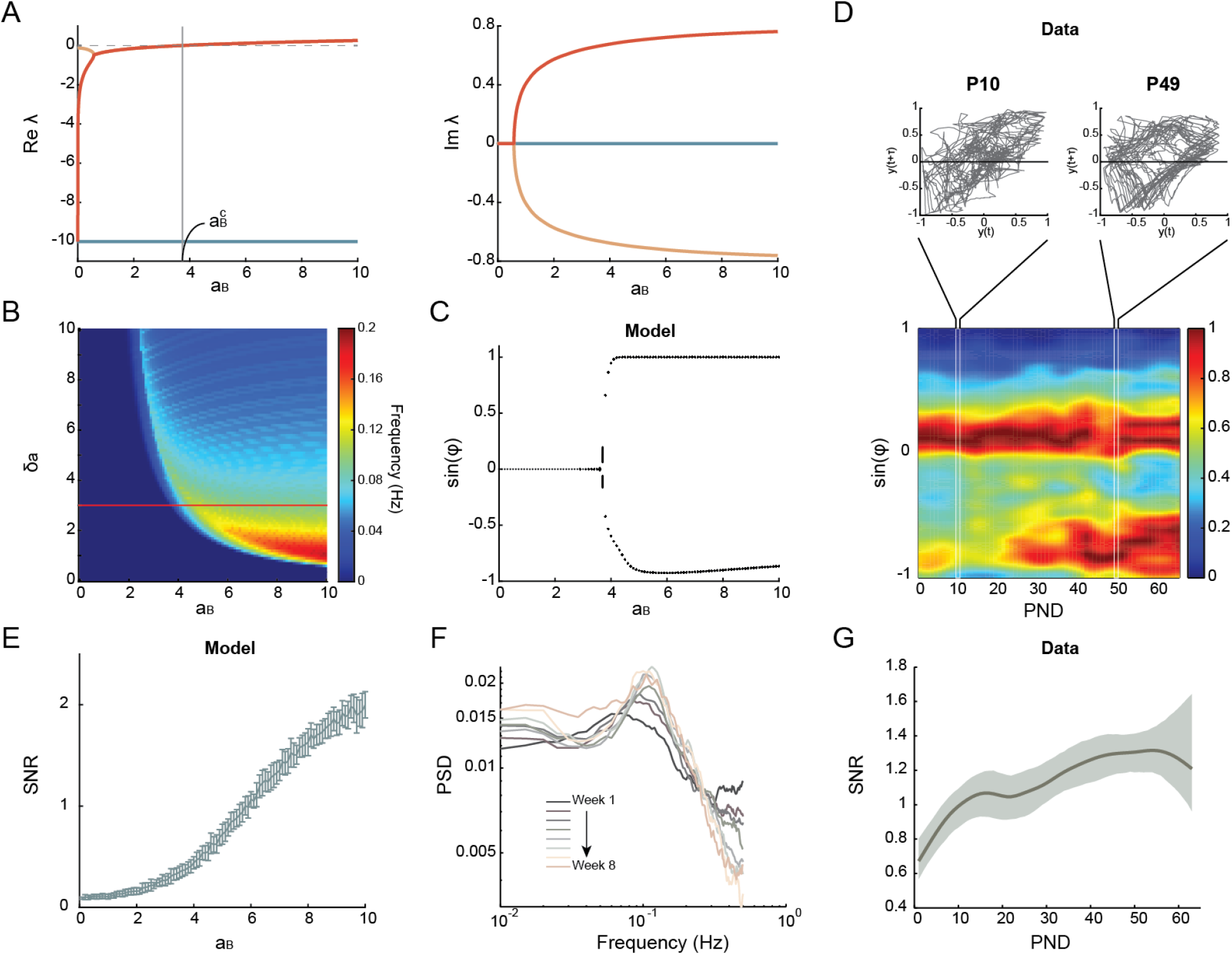
Hopf bifurcation of vocal patterns during development. (A) The real and imaginary parts of the eigenvalues. Jacobian matrix was calculated around the single fixed point (Appendix A). Traces of the same color indicate the real and imaginary parts of the same eigenvalue. 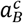 is the critical *a*_*B*_ where a pair of complex conjugate eigenvalues cross the imaginary axis. (B) Frequency map in the *δ*_*a*_ − *a*_*B*_ space. Red line indicates the value used in this study. (C) Phasegram of the deterministic model as a function of *a*_*B*_. (D) Phasegram of data as a function of PND. Color represents probability density normalized per day. Insets are the phase portrait of two examples before and after the bifurcation. (E) Simulated SNR of the stochastic model as a function of *a*_*B*_. Error bars are standard deviations. (F) PSD of vocal amplitude envelope averaged by postnatal weeks. The peak around 0.1 Hz became sharper over time. (G) SNR of data as a function of PND. The curve is fitted by a cubic spline. Shaded area is 95% CI.

The emergence of periodic oscillations can be revealed by a phasegram (Fig. 4C). The phasegram is constructed by the intersection of points between phase portraits under different *a*_*B*_ and a Poincare section [40]. The Poincare section here is a line intersecting the trajectory in the phase space constructed by the sound signal and a delayed version of itself (Methods). When the Hopf bifurcation occurs around 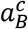, two intersection points instead of one appear, indicating the occurrence of periodic oscillations. Consistent with our model prediction, the phasegram of the vocalization data as a function of postnatal days started exhibiting periodic oscillations after about P30 on average (Fig. 4D). Similar transitions were observed in individual subjects (Fig. S3). Overall, these results support a Hopf bifurcation occurring during vocal development.

The above analyses are based on the deterministic part of the model. When adding the amplitude-dependent Gaussian noise to *v*(*t*), the system is mainly driven by noise at low *a*_*B*_ (Fig. 2B). With *a*_*B*_ above the critical value, the system is dominated by a periodic oscillation (Fig. 2C). With the noise added, the system can occasionally exhibit oscillations at 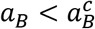, and the closer is *a*_*B*_ to 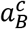, the more frequently oscillations can take place. To quantify the amount of periodic oscillation vs. noise as *a*_*B*_ increases, we calculated the signal-to-noise ratio (SNR). Here we treated the power spectral density (PSD) near 0.1 Hz as signal and PSD at other frequencies as noise. The model predicts that SNR increases with *a*_*B*_ (Fig. 4E). In agreement with this prediction, the PSD of the sound amplitude envelope became more concentrated at 0.1 Hz over time (Fig. 4F), and the SNR increased with postnatal days (p=1.5e-7, MLR; Fig. 4G). Thus, the decreases in the stochasticity and the increases in the regularity of infant vocalization can be explained by the crossing of Hopf bifurcation.

### Interpretations of the control parameter

Arousal allocates energy for behaviors. Contexts inducing higher arousal should have more energy allocated and vice versa. Thus, one natural interpretation for *a*_*B*_ is that it is the baseline of context-dependent energy allocation. Physical development is accompanied by increases in metabolic energy generally, and thus more energy can be allocated for vocalization. During marmoset development, not only did heart rates become higher, but the volume of the heart also expanded. By Fick’s equation, the rate of oxygen consumption is 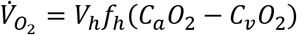, where *V*_*h*_ is the stroke volume of the heart, *f*_*h*_ is the heart rate, and *C*_*a*_*O*_2_ − *C*_*v*_*O*_2_ is the oxygen extraction [41, 42]. Thus, if oxygen extraction in this context is constant across development, then 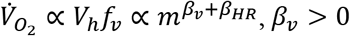. In other words, one should expect the growth of metabolic rate to be faster than heart rate (Fig. 3C). This is consistent with the allometric scaling of *a*_*B*_. The ability to produce regular vocal output thus can be a consequence of general metabolic development.

Previous studies also show that vocal development can be accelerated by increased contingent parental vocal feedback, independent of body growth [10, 43]. In a study of an opposite extreme (lack of parental input), hand-raised marmoset triplets continued producing immature calls and vocal sequences even at one year [44, 45], suggesting that body growth is not a sufficient condition for vocal development. If we decompose the arousal state *a*(*t*) into a growth dependent but socially independent component, e.g., the metabolic energy availability *E*(*t*), and a growth independent but socially tuned factor γ that changes much more slowly: *a*(*t*) = γ*E*(*t*). We can rewrite the last equation of Eq. (1) into

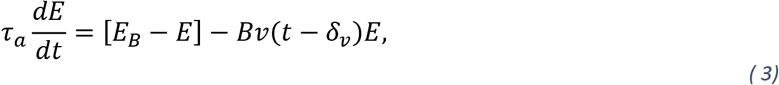

where *E*_*B*_ = *a*_*B*_/γ. *E*_*B*_ can be understood as the baseline metabolic energy, which increases with body growth. The drive *I* can also be rewritten as

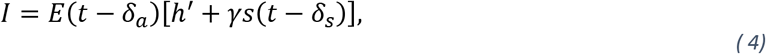

where *h*′ = γ*h*. The socially tuned factor γ is hence simply the strength of the auditory feedback to the vocal production system. In other words, it is the reinforcement strength of the self-produced vocalizations.

An intriguing corollary of this formulation is that the development into the oscillatory vocal output can be accelerated or decelerated by social experiences. For a normally developing individual, *E*_*B*_ increases with body growth, and so the developmental trajectory in the γ – *E*_*B*_ space always goes from left to right (Fig. 5). Increases in *E*_*B*_ also indicate increases in developmental time. Contingent parental feedback can reinforce vocal production which is modeled by the enhancement of γ. Consequently, the trajectory is biased upward in the space (Fig. 5, red lines). With a faster-growing γ, the timing of bifurcation crossing is advanced, and oscillatory vocal output is possible even with a relatively low *E*_*B*_ (Fig. 5 and Fig. S4A). This is consistent with the acceleration of vocal development by contingent parental feedback [10, 43]. Compatible also with empirical findings in hand-raised marmosets, our model shows that, even with a high *E*_*B*_, if the weight of auditory feedback is too weak, the system operates below the bifurcation and thus would still produce immature vocal behavior (Fig. S4B). Therefore, the development of energetics may provide the necessary condition for the animal to produce adult-like vocal behavior, while social feedback provides the correct tuning of it. Both factors may act as control parameters for the development of vocal dynamics.

**Figure 5.**
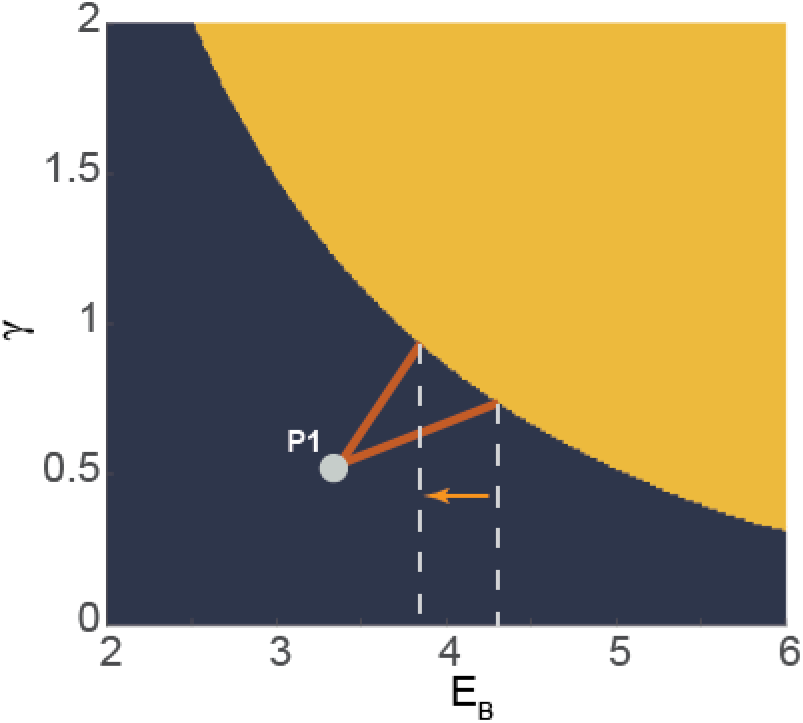
Bifurcation diagram of the deterministic model in the space of *E*_*B*_ and γ. Blue region is the stable fixed point solution, and yellow region is the oscillatory solution. The red line with a steeper slope represents the developmental trajectory with more contingent parental feedback. The arrow indicates the earlier crossing of the bifurcation boundary with more contingent feedback.

## Discussion

In this study, we propose a parsimonious model that integrates sensorimotor processes with the dynamics of internal states based on simultaneous measurements of vocalizations and arousal in infant marmosets. We demonstrated that increases in energy allocation--the baseline arousal level--during development could cause a Hopf bifurcation in vocal sequence production, leading to a stable oscillatory pattern at 0.1 Hz. We further provided insights into how increases in metabolic rate and social feedback can contribute to increasing the value of this parameter. Together, these investigations suggest that motor development can be affected by the development of arousal, which is a quantity for socially guided energy allocation.

Organisms continuously regulate physiological parameters to maintain them within tenable ranges [46]. This is key to survival from the very beginning of life and universal for all species. Arousal is a grand parameter that coordinates physiology at the systems level to accommodate behavioral states to internal demands [47]. Arousal fluctuations correlate with dynamical changes of brain states and behaviors [48-50]. Spontaneously produced motor behaviors are tightly coupled with internal physiological fluctuations, and they are actively coordinated via functionally and structurally interconnected large-scale brain networks [37]. For dedicated brain functions, arousal modulates the gain of visual [51-53] and auditory systems [54], enabling adaptive behaviors [18]. Arousal-modulated sensory processing has been suggested to be regulated via cholinergic input from the ascending arousal system [53, 55]. In the song system of birds, cholinergic modulation of premotor nucleus HVC increases firing rate prior to syllable onsets and enhances song stereotypy, resembling the effect of females’ presence [56]. These changes are consistent with our model's increases in baseline arousal level. Another system implementing feedback loops for motor vigor is the dopaminergic signaling through the basal ganglia, thalamus, and back to the neocortex, which contributes to reward-based motor learning [57, 58]. In birds, the dopamine-secreting neurons in the periaqueductal grey (PAG), which are believed to encode information about social context and arousal, enable vocal learning by releasing dopamine in HVC [59]. This mechanism can facilitate the selection of socially relevant behaviors during development. The neuromodulation for primate and other forms of mammalian vocal development is still unclear and requires future investigation.

Vocal communication is a flexible motor behavior that can be adapted to various contexts through modifying acoustic properties. Our model suggests that drastically changed vocal behavior can be effectively tuned by a slowly varying parameter [60]. This offers a plausible solution for adaptive vocal control. In marmoset monkeys, physical distance consistently induces different vocalizations with shorter, softer, and noisier calls produced in shorter distances [18, 61]. Adult marmosets that are alone (or in an “infinitely far” context) produce the highest arousal levels when compared to other distances between conspecifics [18]. Consistent with our model, animals in this high arousal condition also produce more regular vocalizations [18]. Infant marmosets in the high arousal condition are more likely to reach the limit of their motor and energetic capacity due to a generally high-level motor output. This might explain why we observed a drastic vocal shift in this context during the first two months when the infants were rapidly growing. In comparison, the association between social context and arousal can be a learned trait through experience [10, 43-45]. Without adequate social feedback, the reinforcement of self-produced vocalizations may be downregulated, making the vocal output less vigorous. Social context and arousal are, maybe in this way, naturally linked via the strength of sensory feedback. Future experiments should test the effect of social environment on the strength of auditory feedback and the effect of auditory feedback on the vocal output.

How the system “decides” on the acoustic output could be via the medial brain pathway regulating arousal and motor aspects of vocal production [62-64]. Along the medial wall, several subcortical and cortical areas controlling cardiac activity are related to socially relevant vocal perception and production [35, 37]. The anterior cingulate cortex (ACC), for example, is a robust node along the primate vocal pathway [64, 65]. It reacts strongly to conspecific calls [35] and regulates cardiovascular activity [36, 66, 67]. In the simple scenario depicted in our model, the drive is a conjunctive consequence of sensory input and arousal level. Social context may affect vocal output by shifting the arousal baseline. The integration of social context, arousal, and sensory feedback may occur within ACC. The drive signal can be passed to the downstream motor areas such as PAG, which has been found to initiate vocal production in rodents and primates [68-70]. The PAG dynamics drive the brainstem central pattern generators (CPG) that oscillate on a faster respiratory timescale [60]. The model in this study focuses on the slower dynamics that generate call bouts instead of individual utterances whose boundaries are set by respiration. The acoustic structure is further unpacked via interactions between the CPGs and the vocal apparatus, which changes during development and affects the vocal outcome as well [22]. This study cannot preclude other mechanisms that may produce slow behavioral oscillations, such as the involvement of intrinsic 0.1 Hz fluctuations prevalent in the brain [71]. The biological basis for the long-delayed arousal signal giving rise to this temporal scale requires future studies to verify.

Energy is often an ignored factor when it comes to motor control. Energy impacts motor output at various timescales. Sound production, for instance, is a prerequisite for self-preservation in some species; its importance is reflected in the generally high energetic cost attained by a variety of physiological and biomechanical mechanisms [72]. In some arthropods and frogs, the cost of calling can be up to 20 times the resting metabolic rate [73]. At short timescales, this dependence on energetics affects acoustic features such as power, frequency, duration, and rate of call emission [74]. At longer timescales, the internal goals and needs of the animal change, and energy allocated for sound production varies according to the time of the day [75], nutrition [76], seasons [77], state of learning [78], and age [79]. In our study here, we offered an energetic interpretation for our model to raise attention to this critical factor. There are several mechanisms by which metabolic energy affects neural activity. At the cellular and tissue levels, energy availability can influence neural excitability and information processing [80, 81]. At the whole-body level, diet, exercise, and blood sugar levels affect neural signaling [82]. The effect of metabolism on behavioral phenotypes has been linked to biochemical pathways operating in the brain. For example, in insects, inhibition of oxidative phosphorylation (a process producing maximal ATP) causes increased aggression [83]. Moreover, the effect of oxidative phosphorylation inhibition is modulated by the social environment [83], suggesting the context-dependent energy demand for producing motor behavior. Future studies bridging energetics and behavior at the levels of genetics, biochemistry, neurons, and physiology will help fill in the details of our theory.

## Material and methods

### Subjects

The subjects used in the original study were 17 infants up to two months of age, captive common marmosets (*Callithrix jacchus*) housed at Princeton University. Vocalization recordings from ten infants have been published previously [10]. Another seven infants were used for heart rate measurements throughout their development, and the data were used in a previous study [25]. All experiments were performed in compliance with the guidelines of the Princeton University Institutional Animal Care and Use Committee.

### Vocalization recordings

Ten infant marmosets participated in this experiment. On each day starting from the first postnatal day, an infant was temporarily separated from its parents and taken to the experiment room, where it was placed in a test box. Undirected vocalizations were recorded for 5 min per day. The types of vocalizations that were observed were identical to those produced when the infant was naturally separated from its parents. A speaker was placed in the room broadcasting pink noise (amplitude decaying inversely proportional to frequency) of 45 dB (at 0.88m from the speaker to the microphone) in order to mask occasional noises produced outside of the testing room. The sound pressure level (SPL) was measured at the microphone location and used as a reference to calibrate the SPL of marmoset vocalizations. The digital recorder (ZOOM H4n Handy Recorder) was placed directly in front of the transfer cage at a distance of 0.76m. Audio signals were acquired at a sampling frequency of 96kHz.

### Electrocardiogram (ECG) recording and heart rate analysis

A different group of infant marmosets (n=7) participated in this experiment. Infant marmosets were placed in a test box and recorded as they were spontaneously vocalizing for 10 mins. To record the electrocardiogram (ECG) signal, we used Ag-AgCl surface electrodes (Grass Technology) placed on the chest, close to the heart. The electrodes were sewn onto an elastic band that was secured around the marmoset’s thorax. The ECG signals were acquired by the OmniPlex data acquisition system (Plexon) at a sample rate of 40 kHz. The ECG signals were downsampled and bandpass-filtered between 5 and 39 Hz. Heartbeats were detected using template matching by cross-correlation within a 50 ms sliding window. The heartbeats were binarized and convolved with a Gaussian window with σ = 0.25 s. The mean heart rate of the session was estimated as the mean of this convolved time series.

### Acoustic feature processing

To calculate the acoustic power of marmoset vocalizations, we removed non-vocal signals by identifying the boundaries of individual utterances. Vocal utterances were defined as continuous sound separated by silence. The boundaries of utterances were determined by threshold crossings of the sound amplitude envelope and checked manually. Sound pressure was calculated as the root-mean-square (RMS) envelope of the sound signal and calibrated with the 45-dB pink noise background. It was then converted to power in milliwatts using a standard equation that accounts for measurement distance (0.76m) and assumes hemispherical spreading 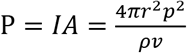, where *p* the RMS sound pressure in Pa, *ρ*=1.2 kg m^-3^ the air density, and *v*=344 m s^-1^ the speed of sound [84].

### Spectral analysis

The acoustic power of the sound amplitude envelope was downsampled to 1 Hz. The PSD of was calculated using MATLAB implementation of the multi-taper method *pmtm*. Mean PSDs were calculated for every postnatal week up to the 8^th^ week across all ten marmosets. The SNR was estimated as the ratio between the sum of PSD within the range of 0.05 – 0.2 Hz and the sum of the rest within 0.02 – 0.5 Hz [85].

### Heart rate analysis

To analyze the heart rate dynamics around vocal production, we first converted detrended momentary heart rates into percentiles. This was done by binning 100 percentiles of the session heart rate and assigning the heart rate of each time point into a percentile bin.

To correlate heart rate drop with acoustic power, we calculated the heart rate change from call onset to the lowest heart rate during call production. The acoustic power was standardized by session, and the mean power of a call was calculated for the correlation.

### Model setup

We modeled the response function of the arousal gained auditory feedback to the neural population as a sigmoidal curve. We added noise 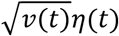 to the deterministic system, which is a delta-correlated Gaussian white noise with zero mean and an activity-dependent amplitude. The square-root scaling is due to the central limit theorem, as we assume that the activity of each neuron in this population is an independent stochastic process [86]. We only introduced noise in the modeling of neural activity for simplicity; in principle, there is also noise in arousal and vocalization.

The stochastic differential equation with multiplicative noise was solved in two steps. The stochastic part was integrated first to generate a sample of neural activity *v*^∗^ with guaranteed positivity, using a trick developed in ref. [87]. In the next step, *v*^∗^ was used as a starting point for the integration of the deterministic part. The time step was set at dt=0.01.

### Parameter estimation

To estimate *a*_*B*_ of the model, we performed an unscented Kalman filter for noisy data to track observations of vocalization and heart rate [39]. We also included covariance inflation due to the noisiness of the data [88]. The remaining parameters were fixed according to Table 1. For each session, we normalized the sound amplitude by the maximum. We detrended the heart rate signal, band-passed it between 0.02 and 1 Hz, and converted it to percentiles. We then scaled the signal from 1 to 10 and passed each scaled version through the Kalman filter tracking. To find the best scaling factor, we calculated the tracking error of sound and heart rate as the sum of the L2 norm of these two errors and selected the scaling factor yielding the lowest error. We initialized the covariance matrix for real-time tracking of state variables using the sample covariance matrix of sound and heart rate signals. The process noise for parameter tracking was set to 0.015 and was reset each iteration. The time-series data were repeated for each session such that enough iterations were given for stabilizing the tracking. The estimated *a*_*B*_ was calculated as the mean of the second half of the tracking.

### Bifurcation analysis of data

To construct the phasegram of vocalization data, we first calculate the envelope of the acoustic power using the RMS with a 1-s sliding window. In this way, we smooth out the rhythms at the respiratory timescale. We then Hilber-transformed the signal and estimated the time-dependent phase angle *ϕ*(*t*). This step mitigated the observational variability of acoustic power. We finally took *y*(*t*) = sin (*ϕ*(*t*)) to construct a phase portrait in the space of *y*(*t*) and *y*(*t* + τ), where τ is a time delay. τ is typically one-tenth to one-half the oscillation period [89], and in this analysis, we used τ = 1.5 s. The Poincare section was chosen as the x-axis of the phase space. We then calculated a Gaussian-filtered 2D histogram using MATLAB function *ndhist* (file exchange #45325) in the space of postnatal days (PND) and sin (*ϕ*). The histogram was normalized within the same postnatal day. A similar procedure was adopted to calculate the simulated data.

### Statistical analysis

We used the MATLAB *fitlm* routine to fit robust linear models: *y* = *b*_0_ + *b*_1_*x* + *b*_2_*ID* + *∈*, where *y* is the response, *x* is the independent variable, and *ID* is the subject, to find the scaling exponent of heart rate vs. body mass and *a*_*B*_ vs. body mass. We assumed that each individual had its own slope and intercept and thus included the identities of the subjects as a predictor in the model. A similar linear fitting was applied to test the dependence of *a*_*B*_ and SNR on PND. For linear fitting of heart rate drop vs. acoustic power, we also included PND as a predictor in the MLR model. To compare the allometric scaling of heart rate vs. *a*_*B*_, we fitted the robust multiple linear model log(*a*_*B*_) − log(*hr*) = *b*_0_ + *b*_1_ log(*weight*) + *b*_2_*ID* + *∈* and tested if *b*_1_ was significantly different from zero. The statistical significance was evaluated based on the p-values reported in the MATLAB function output.

The test for significantly high or low heart rate was performed by randomizing the physiological signals and calculating the mean 1000 times to construct the 95% CI of the null distribution. The cross correlation between sound amplitude and heart rate was calculated by sessions and tested against 0 using t-test with Bonferroni correction for each lag point. We adopted the Type I error α = 0.05 for the criterion of statistical significance.

## Author contributions

**Yisi Zhang:** Conceptualization, Methodology, Software, Validation, Formal analysis, Data Curation, Writing – Original Draft, Writing – Review & Editing, Visualization. **John Alvarez:** Formal analysis, Data Curation. **Asif Ghazanfar:** Conceptualization, Writing – Original Draft, Writing – Review & Editing, Supervision, Funding acquisition.

## Acknowledgment

We thank Daniel Takahashi for helpful discussions. This research was supported by the NIH NINDS (R01NS054898) to AAG.

## Appendix A. Derivation of bifurcation

The system has fixed points satisfying

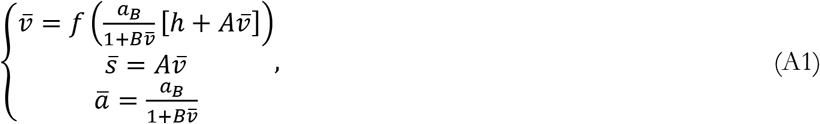

where 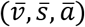 is the fixed point, *A*, *B*, *a*_*B*_, *h* > 0. The equations can be rearranged into solving a cubic equation in terms of 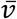

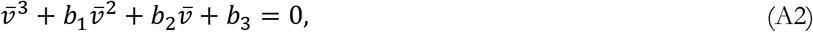

where 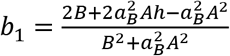, 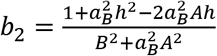, and 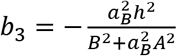. Since *b*_3_ < 0, there are either 1 or 3 positive real roots. Furthermore, when *a*_*B*_ → 0^+^, it is easy to show that the single positive root 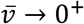, and *b*_3_ is a monotonic decreasing function of *a*_*B*_. Thus, there is always a single positive fixed point in this system.

Oscillations may occur where in parameter space this fixed point becomes unstable. To investigate the stability of the fixed point, we linearize the system around it by writing

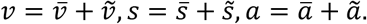

For convenience, we neglect the tilde and still use *v*, *s*, *a* in the following but for perturbations around the fixed point. The linearized system is

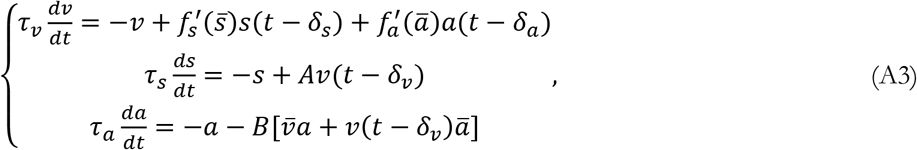

where 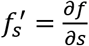 and 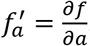.

Now we look for solutions in the form *v* = *v*_0_*e*^λ*t*^, *s* = *s*_0_*e*^λ*t*^ and *a* = *a*_0_*e*^λ*t*^. Plugging into the linearized equations gives

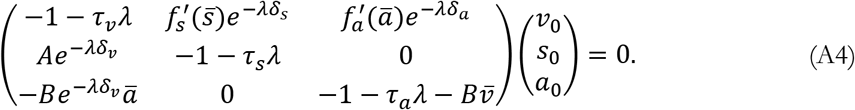

λ thus must be a root of the transcendental equation

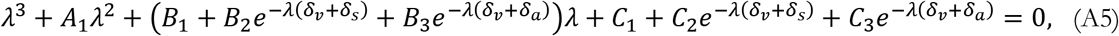

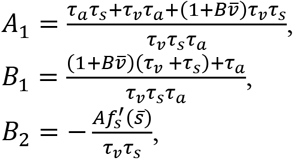

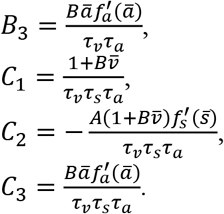

First, we check the stability with the delays *δ*_*v*_ = *δ*_*s*_ = *δ*_*a*_ = 0, and (A5) becomes

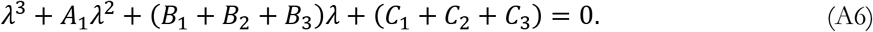

We make the following hypotheses

(H1) *Bh* ≥ *A*.

(H2) *A*_1_(*B*_1_ + *B*_2_ + *B*_3_) > *C*_1_ + *C*_2_ + *C*_3_.

(H1) sufficiently guarantees *C*_1_ + *C*_2_ + *C*_3_ > 0. It is also easy to check that for *a*_*B*_ → 0^+^ as well as for *a* → +∞, (H2) is met, and so a wide range of parameters ensures (H2) in *a*_*B*_ ∈ (0, ∞). With *A*_1_ > 0, (H1) and (H2), the Routh-Hurwitz criterion is satisfied to have *Re* λ < 0. Thus, without delay, 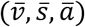 is asymptotically stable.

The delay-destabilized systems have been widely studied, and here, we suppose some delay exists for the system to exhibit oscillations. We further assume that *δ*_*a*_ ≫ *δ*_*v*_, *δ*_*s*_, and in the following analyses, we approximate *δ*_*v*_, *δ*_*s*_ ≈ 0 without loss of generality. Substituting λ = *μ* + *iω* into the characteristic equation (A5) and for the real and imaginary parts we have

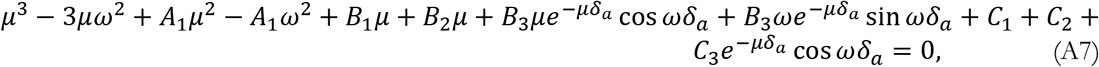

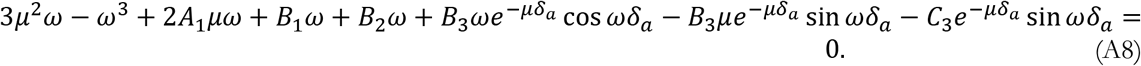

Now suppose for some *δ*_*a*_, (A5) has purely imaginary solutions λ = ±*iω*. At *μ* = 0, we have

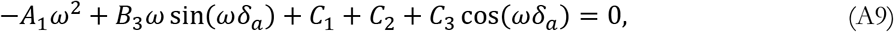

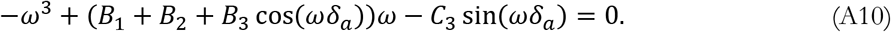

Let 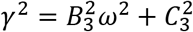 and 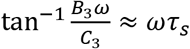 for *ω*τ_*s*_ ≪ 1, (A9) and (A10) becomes

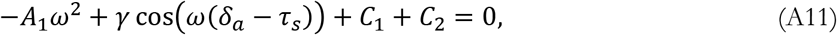

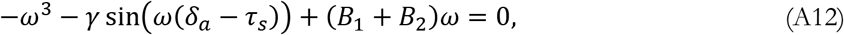

from which we get

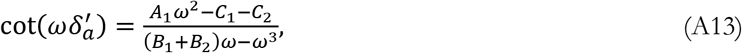

where 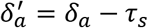. Suppose 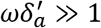, the smallest *ω* = *ω*_0_ that solves (A13) is when 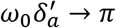.

Expanding (A13) around here follows

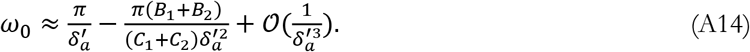

One can show that for 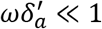, a positive solution for *ω*_0_ does not exist. Thus, only when 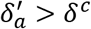, we have *ω*_0_ > 0. Now suppose this condition is met, squaring and adding (A11) and (A12), one gets

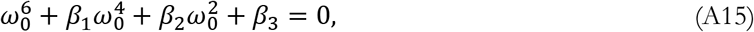

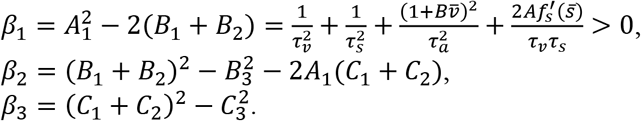

For *a*_*B*_ → 0^+^, we have *β*_2_, *β*_3_ > 0, *β*_1_*β*_2_ > *β*_3_, and there are no positive roots for *ω*^2^. In this situation, there is only *μ* < 0. As *a*_*B*_ increases, if there is an 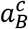 at which (A15) starts to hold with positive *ω*^2^, *μ* = 0 starts to exist. In other words, the critical 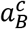 should occur when 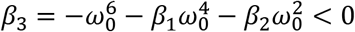. Since *C*_1_ + *C*_2_+ *C*_3_ > 0, this is essentially at 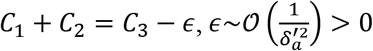. Ignoring *∈*, it approximately corresponds to the positive roots of the following equation

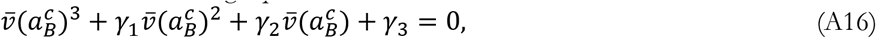

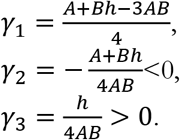

Here 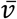 is a monotonic increasing function of *a*_*B*_ (as 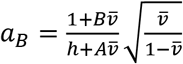 a monotonic increasing function of 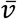 for *Bh* > *A*). The signs of the coefficients produce two positive roots for 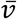, indicating that *β*_3_ starts from positive (when *a*_*B*_ → 0^+^), decreases to negative and comes back to positive again as *a*_*B*_ increases from 0 to +∞. Thus, there is a range *a*_*B*_ ∈ (*a*_*B*1_, *a*_*B*2_) with the boundaries approximately satisfying (A16) produces oscillations.

From (A5), it can be demonstrated that at λ = ±*iω*_0_, 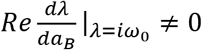. Thus, there are two critical 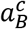 where Hopf bifurcation occurs. Outside this regime, no real solution for *ω* exists, i.e., *μ* = 0 cannot be satisfied. With *a*_*B*_ < *a*_*B*1_, the system remains stable like the case of the immature phase of vocal development; with *a*_*B*_ > *a*_*B*2_, the system is saturated and becomes stable again. The second case is considered physiologically infeasible in this study.

## Appendix B. Derivation of period

From (A14), it is easy to get

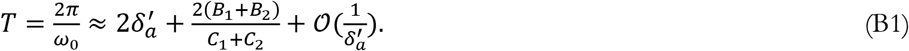

The oscillation period *T* is thus approximately twice of 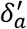 plus an *a_B_* dependent term. Note that for 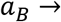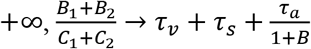. Thus, with *B* ≫ 1 and τ_*v*_, τ_*s*_ ≪ *δ_a_*, when *a_B_* is sufficiently large, the period is approximately 2*δ*_*a*_.

## Supplementary Figures

**Figure S1.**
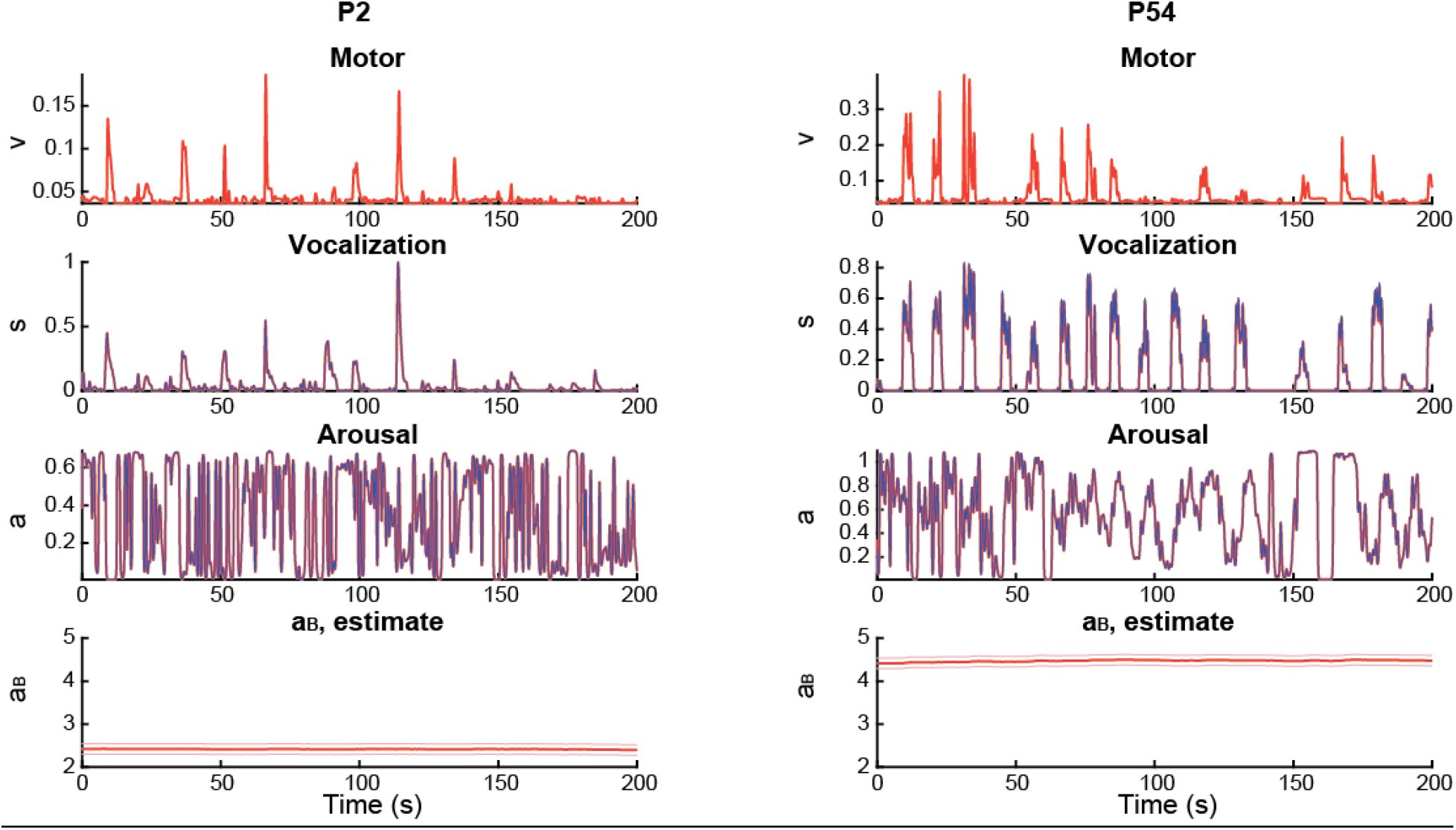
Kalman filter tracking of vocalization (*s*(*t*)) and heart rate (*a*(*t*)) and estimation of *a*_*B*_. Red traces are state tracking, blue traces are observation. *a*_*B*_ errors are predetermined process noise.

**Figure S2.**
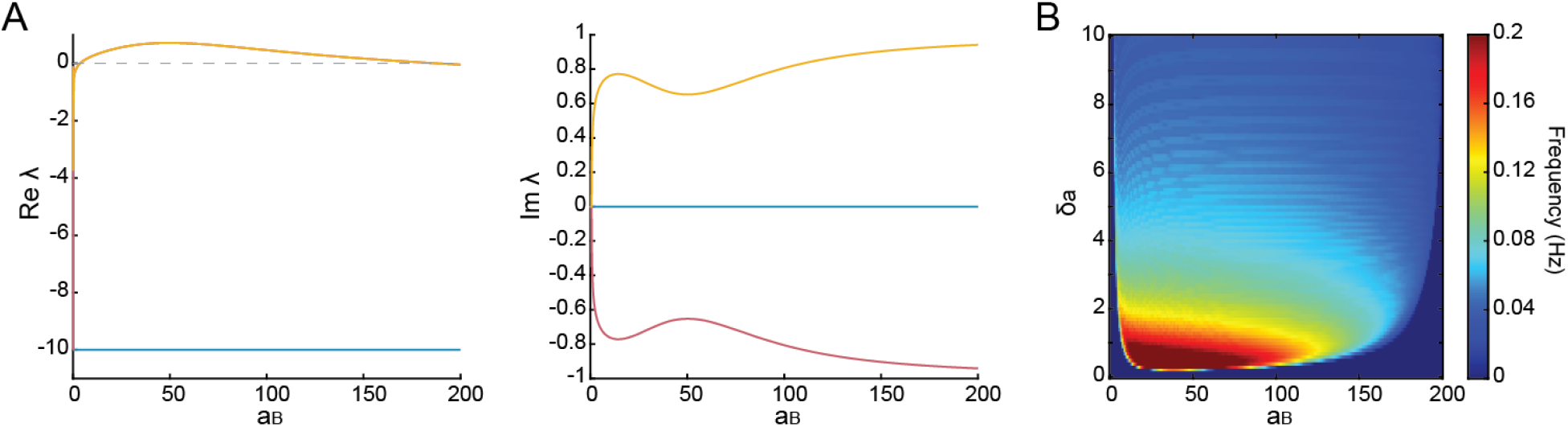
Hopf bifurcation at larger *a*_*B*_. (A) The real (left) and imaginary (right) parts of the eigenvalue λ. Two zero-crossing points exist for *Re* λ in *a*_*B*_ ∈ (0, +∞). The interval with positive *Re* λ produces oscillations. (B) The frequency map in the *δ*_*a*_ - *a*_*B*_ space.

**Figure S3.**
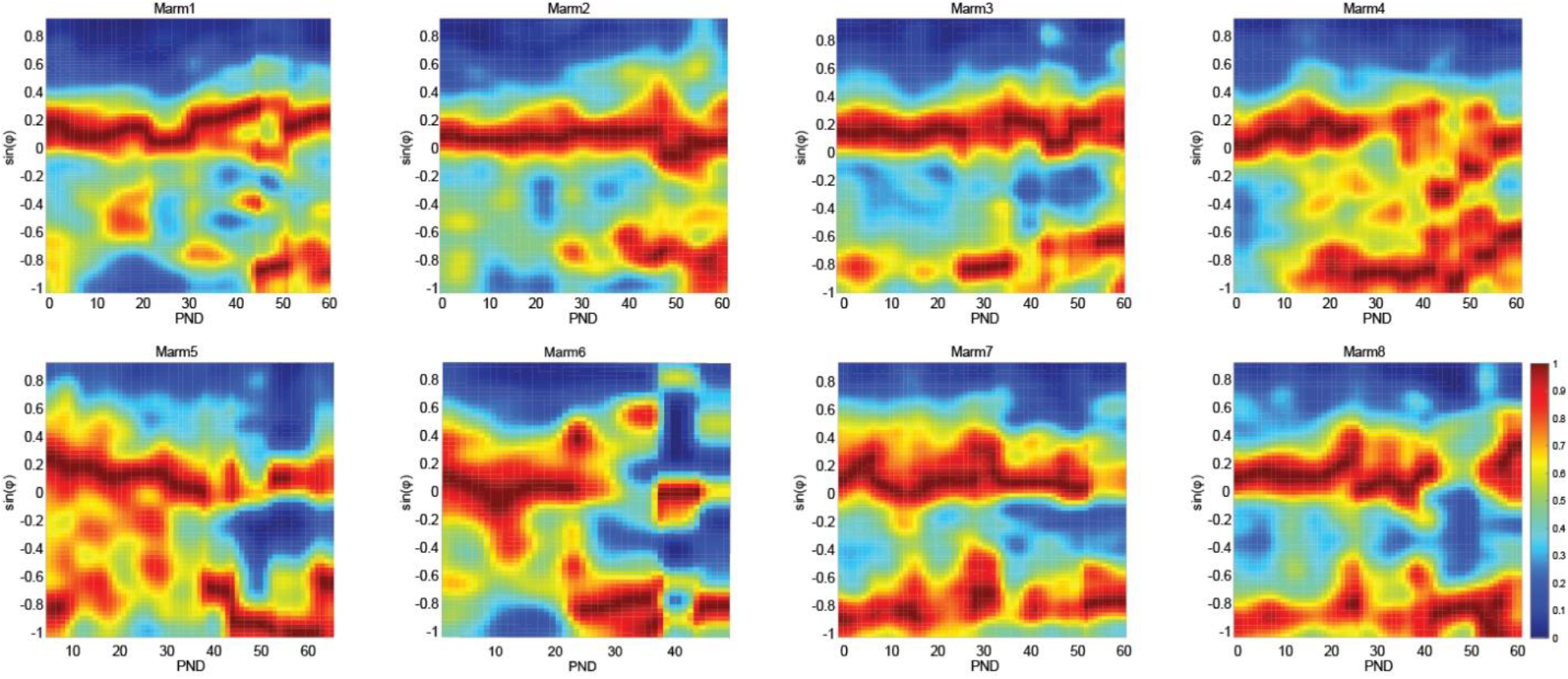
Phasegram of individuals. These are 8 out of 10 individuals, the other 2 individuals have too few time points to plot. The last two subjects already started producing periodic oscillatory vocalizations from P1.

**Figure S4.**
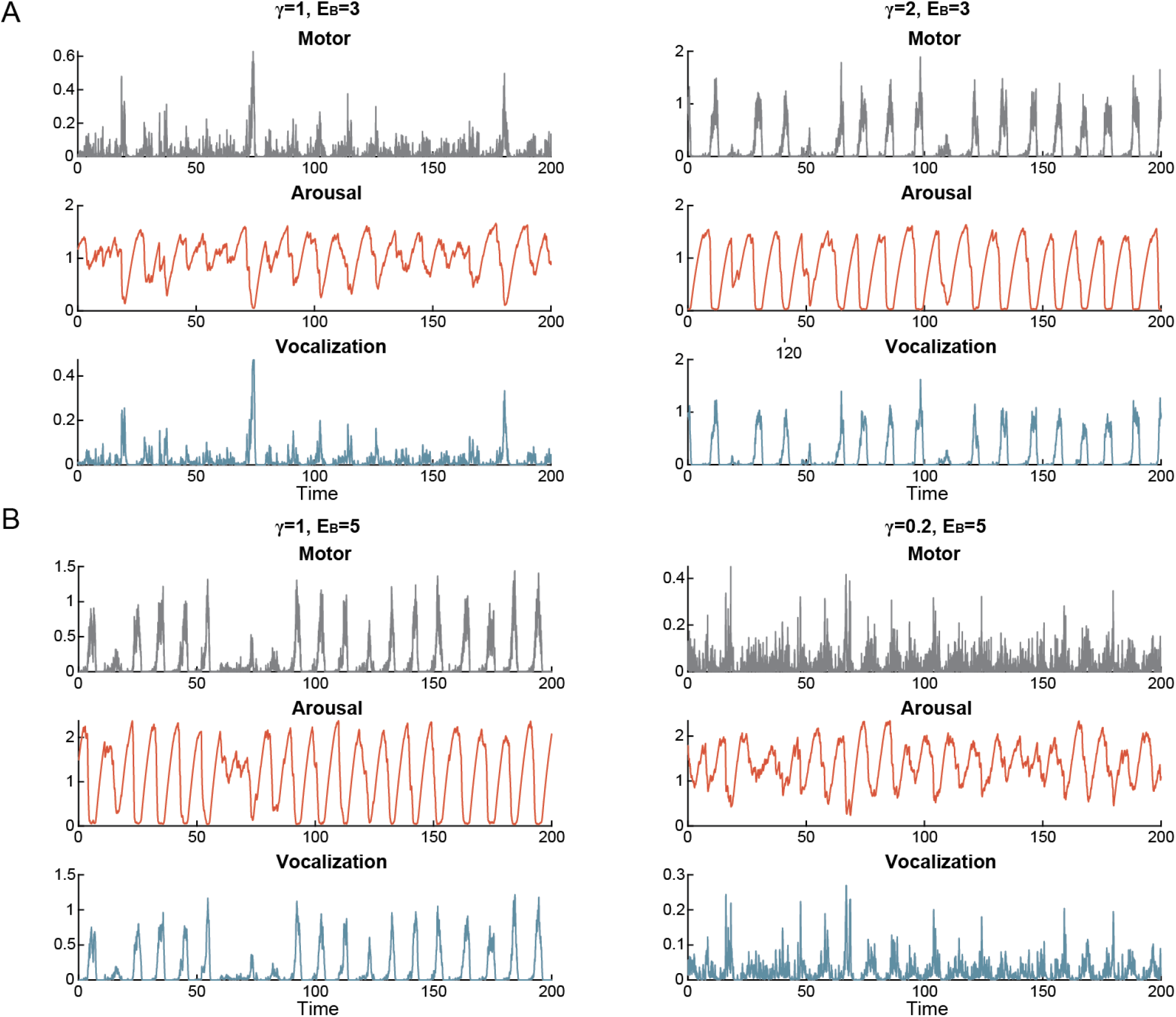
Acceleration and deceleration of vocal development by tuning the social factor γ. (A) High γ can cause oscillation even with low *E*_*B*_. Left: an example of low *E*_*B*_ and moderate γ. Right: an example of low *E*_*B*_ and high γ. (B) Low γ can impede the development of oscillatory pattern even with high *E*_*B*_. Left: an example of high *E*_*B*_ and moderate γ. Right: an example of high *E*_*B*_ and low γ.

## References

1. Lummaa V. Why Cry? Adaptive Significance of Intensive Crying in Human Infants. Evolution and Human Behavior. 1998;19(3):193–202. doi: 10.1016/s1090-5138(98)00014-2.

2. Newman JD. The infant cry of primates. Infant crying: Springer; 1985. p. 307–23.

3. Kent RD, Murray AD. Acoustic features of infant vocalic utterances at 3, 6, and 9 months. The Journal of the Acoustical Society of America. 1982;72(2):353–65. PubMed PMID: 7119278.

4. Scheiner E, Hammerschmidt K, Jürgens U, Zwirner P. Acoustic Analyses of Developmental Changes and Emotional Expression in the Preverbal Vocalizations of Infants. Journal of Voice. 2002;16(4):509–29. doi: 10.1016/s0892-1997(02)00127-3.

5. Lipkind D, Marcus GF, Bemis DK, Sasahara K, Jacoby N, Takahasi M, et al. Stepwise acquisition of vocal combinatorial capacity in songbirds and human infants. Nature. 2013;498(7452):104–8. doi: 10.1038/nature12173. PubMed PMID: 23719373; PubMed Central PMCID: PMC3676428.

6. Sasahara K, Tchernichovski O, Takahasi M, Suzuki K, Okanoya K. A rhythm landscape approach to the developmental dynamics of birdsong. Journal of the Royal Society, Interface / the Royal Society. 2015;12(112). doi: 10.1098/rsif.2015.0802. PubMed PMID: 26538559; PubMed Central PMCID: PMC4685852.

7. Prat Y, Taub M, Yovel Y. Vocal learning in a social mammal: Demonstrated by isolation and playback experiments in bats. Science Advances. 2015;1(2):e1500019. doi: 10.1126/sciadv.1500019.

8. Zhang YS, Ghazanfar AA. Perinatally Influenced Autonomic System Fluctuations Drive Infant Vocal Sequences. Current biology : CB. 2016. doi: 10.1016/j.cub.2016.03.023. PubMed PMID: 27068420.

9. Gultekin YB, Hildebrand DGC, Hammerschmidt K, Hage SR. High plasticity in marmoset monkey vocal development from infancy to adulthood. Science Advances. 2021;7(27). doi: 10.1126/sciadv.abf2938.

10. Takahashi DY, Fenley AR, Teramoto Y, Narayanan DZ, Borjon JI, Holmes P, et al. The developmental dynamics of marmoset monkey vocal production. Science. 2015;349(6249):734–8. doi: 10.1126/science.aab1058.

11. Teramoto Y, Takahashi DY, Holmes P, Ghazanfar AA. Vocal development in a Waddington landscape. eLife. 2017;6. doi: 10.7554/eLife.20782.

12. Borjon JI, Takahashi DY, Cervantes DC, Ghazanfar AA. Arousal dynamics drive vocal production in marmoset monkeys. Journal of neurophysiology. 2016;116(2):753–64. doi: 10.1152/jn.00136.2016.

13. Schehka S, Esser K-H, Zimmermann E. Acoustical expression of arousal in conflict situations in tree shrews (Tupaia belangeri). Journal of Comparative Physiology A. 2007;193(8):845–52. doi: 10.1007/s00359-007-0236-8.

14. Rendall D. Acoustic correlates of caller identity and affect intensity in the vowel-like grunt vocalizations of baboons. The Journal of the Acoustical Society of America. 2003;113(6):3390. doi: 10.1121/1.1568942.

15. Cooper BG, Goller F. Physiological Insights Into the Social-Context-Dependent Changes in the Rhythm of the Song Motor Program. Journal of neurophysiology. 2006;95(6):3798–809. doi: 10.1152/jn.01123.2005.

16. Kao MH, Doupe AJ, Brainard MS. Contributions of an avian basal ganglia–forebrain circuit to real-time modulation of song. Nature. 2005;433(7026):638–43. doi: 10.1038/nature03127.

17. PFAFF DW. Brain Arousal and Information Theory: Neural and Genetic Mechanisms: Harvard University Press; 2006.

18. Liao DA, Zhang YS, Cai LX, Ghazanfar AA. Internal states and extrinsic factors both determine monkey vocal production. Proceedings of the National Academy of Sciences. 2018;115(15):3978–83. doi: 10.1073/pnas.1722426115.

19. Hessler NA, Doupe AJ. Social context modulates singing-related neural activity in the songbird forebrain. Nature Neuroscience. 1999;2(3):209–11. doi: 10.1038/6306.

20. Thelen E, Smith LB. A dynamic systems approach to the development of cognition and action: MIT press; 1996.

21. Smith LB, Thelen E. Development as a dynamic system. Trends in cognitive sciences. 2003;7(8):343–8. doi: 10.1016/s1364-6613(03)00156-6.

22. Zhang YS, Ghazanfar AA. Vocal development through morphological computation. PLoS biology. 2018;16(2). doi: 10.1371/journal.pbio.2003933.

23. Fogel A, Thelen E. Development of early expressive and communicative action: Reinterpreting the evidence from a dynamic systems perspective. Developmental Psychology. 1987;23(6):747–61. doi: 10.1037/0012-1649.23.6.747.

24. Zhang YS, Takahashi DY, Liao DA, Ghazanfar AA, Elemans CPH. Vocal state change through laryngeal development. Nature communications. 2019;10(1). doi: 10.1038/s41467-019-12588-6.

25. Gustison ML, Borjon JI, Takahashi DY, Ghazanfar AA. Vocal and locomotor coordination develops in association with the autonomic nervous system. eLife. 2019;8. doi: 10.7554/eLife.41853.

26. Blumberg Mark S, Coleman Cassandra M, Gerth Ashlynn I, McMurray B. Spatiotemporal Structure of REM Sleep Twitching Reveals Developmental Origins of Motor Synergies. Current Biology. 2013;23(21):2100–9. doi: 10.1016/j.cub.2013.08.055.

27. Petersson P, Waldenström A, Fåhraeus C, Schouenborg J. Spontaneous muscle twitches during sleep guide spinal self-organization. Nature. 2003;424(6944):72–5. doi: 10.1038/nature01719.

28. Khazipov R, Sirota A, Leinekugel X, Holmes GL, Ben-Ari Y, Buzsáki G. Early motor activity drives spindle bursts in the developing somatosensory cortex. Nature. 2004;432(7018):758–61. doi: 10.1038/nature03132.

29. Tiriac A, Uitermarkt Brandt D, Fanning Alexander S, Sokoloff G, Blumberg Mark S. Rapid Whisker Movements in Sleeping Newborn Rats. Current Biology. 2012;22(21):2075–80. doi: 10.1016/j.cub.2012.09.009.

30. Blumberg Mark S, Marques Hugo G, Iida F. Twitching in Sensorimotor Development from Sleeping Rats to Robots. Current Biology. 2013;23(12):R532–R7. doi: 10.1016/j.cub.2013.04.075.

31. Moulin-Frier C, Nguyen SM, Oudeyer P-Y. Self-organization of early vocal development in infants and machines: the role of intrinsic motivation. Frontiers in Psychology. 2014;4. doi: 10.3389/fpsyg.2013.01006.

32. Warlaumont AS, Westermann G, Buder EH, Oller DK. Prespeech motor learning in a neural network using reinforcement. Neural Networks. 2013;38:64–75. doi: 10.1016/j.neunet.2012.11.012.

33. Wittenbach JD, Bouchard KE, Brainard MS, Jin DZ. An Adapting Auditory-motor Feedback Loop Can Contribute to Generating Vocal Repetition. Plos Comput Biol. 2015;11(10):e1004471. doi: 10.1371/journal.pcbi.1004471.

34. Lorenz K. The foundations of ethology: Springer verlag; 1981.

35. Takahashi DY, El Hady A, Zhang YS, Liao DA, Montaldo G, Urban A, et al. Social-vocal brain networks in a non-human primate. 2021. doi: 10.1101/2021.12.01.470701.

36. Alexander L, Clarke H, Roberts A. A Focus on the Functions of Area 25. Brain Sciences. 2019;9(6):129. doi: 10.3390/brainsci9060129.

37. Zhang YS, Takahashi DY, El Hady A, Liao DA, Ghazanfar AA. Active neural coordination of motor behaviors with internal states. bioRxiv. 2021:2021.12.10.472142. doi: 10.1101/2021.12.10.472142.

38. Enoka RM, Duchateau J. Rate Coding and the Control of Muscle Force. Cold Spring Harbor Perspectives in Medicine. 2017;7(10):a029702. doi: 10.1101/cshperspect.a029702.

39. Voss HU, Timmer J, Kurths J. Nonlinear Dynamical System Identification from Uncertain and Indirect Measurements. International Journal of Bifurcation and Chaos. 2011;14(06):1905–33. doi: 10.1142/s0218127404010345.

40. Herbst CT, Herzel H, Švec JG, Wyman MT, Fitch TW. Visualization of system dynamics using phasegrams. Journal of The Royal Society Interface. 2013;10(85):20130288. doi: 10.1098/rsif.2013.0288.

41. White CR, Kearney MR. Metabolic Scaling in Animals: Methods, Empirical Results, and Theoretical Explanations. 2014:231–56. doi: 10.1002/cphy.c110049.

42. Fick A. Ueber die Messung des Blutquantum in den Herzventrikeln. Sb Phys Med Ges Worzburg. 1870:16–7.

43. Takahashi DY, Liao DA, Ghazanfar AA. Vocal Learning via Social Reinforcement by Infant Marmoset Monkeys. Current biology : CB. 2017;27(12):1844–52 e6. doi: 10.1016/j.cub.2017.05.004. PubMed PMID: 28552359.

44. Gultekin YB, Hage SR. Limiting parental feedback disrupts vocal development in marmoset monkeys. Nature communications. 2017;8:14046. doi: 10.1038/ncomms14046.

45. Gultekin YB, Hage SR. Limiting parental interaction during vocal development affects acoustic call structure in marmoset monkeys. Science Advances. 2018;4(4). doi: 10.1126/sciadv.aar4012.

46. Cannon WB. Organization for Physiological Homeostasis. Physiological reviews. 1929;9(3):399–431. doi: 10.1152/physrev.1929.9.3.399.

47. Sterling P. Allostasis: A model of predictive regulation. Physiology & behavior. 2012;106(1):5–15. doi: 10.1016/j.physbeh.2011.06.004.

48. McGinley Matthew J, Vinck M, Reimer J, Batista-Brito R, Zagha E, Cadwell Cathryn R, et al. Waking State: Rapid Variations Modulate Neural and Behavioral Responses. Neuron. 2015;87(6):1143–61. doi: 10.1016/j.neuron.2015.09.012.

49. Raut RV, Snyder AZ, Mitra A, Yellin D, Fujii N, Malach R, et al. Global waves synchronize the brain’s functional systems with fluctuating arousal. Science Advances. 2021;7(30):eabf2709. doi: 10.1126/sciadv.abf2709.

50. Fontanini A, Katz DB. Behavioral States, Network States, and Sensory Response Variability. Journal of neurophysiology. 2008;100(3):1160–8. doi: 10.1152/jn.90592.2008.

51. Niell CM, Stryker MP. Modulation of Visual Responses by Behavioral State in Mouse Visual Cortex. Neuron. 2010;65(4):472–9. doi: 10.1016/j.neuron.2010.01.033.

52. Fu Y, Tucciarone Jason M, Espinosa JS, Sheng N, Darcy Daniel P, Nicoll Roger A, et al. A Cortical Circuit for Gain Control by Behavioral State. Cell. 2014;156(6):1139–52. doi: 10.1016/j.cell.2014.01.050.

53. Pinto L, Goard MJ, Estandian D, Xu M, Kwan AC, Lee S-H, et al. Fast modulation of visual perception by basal forebrain cholinergic neurons. Nature Neuroscience. 2013;16(12):1857–63. doi: 10.1038/nn.3552.

54. McGinley Matthew J, David Stephen V, McCormick David A. Cortical Membrane Potential Signature of Optimal States for Sensory Signal Detection. Neuron. 2015;87(1):179–92. doi: 10.1016/j.neuron.2015.05.038.

55. Reimer J, McGinley MJ, Liu Y, Rodenkirch C, Wang Q, McCormick DA, et al. Pupil fluctuations track rapid changes in adrenergic and cholinergic activity in cortex. Nature communications. 2016;7(1). doi: 10.1038/ncomms13289.

56. Jaffe PI, Brainard MS. Acetylcholine acts on songbird premotor circuitry to invigorate vocal output. eLife. 2020;9. doi: 10.7554/eLife.53288.

57. Turner RS, Desmurget M. Basal ganglia contributions to motor control: a vigorous tutor. Current opinion in neurobiology. 2010;20(6):704–16. doi: 10.1016/j.conb.2010.08.022.

58. Dudman JT, Krakauer JW. The basal ganglia: from motor commands to the control of vigor. Current opinion in neurobiology. 2016;37:158–66. doi: 10.1016/j.conb.2016.02.005.

59. Tanaka M, Sun F, Li Y, Mooney R. A mesocortical dopamine circuit enables the cultural transmission of vocal behaviour. Nature. 2018;563(7729):117–20. doi: 10.1038/s41586-018-0636-7.

60. Zhang YS, Ghazanfar AA. A Hierarchy of Autonomous Systems for Vocal Production. Trends in neurosciences. 2020;43(2):115–26. doi: 10.1016/j.tins.2019.12.006.

61. Choi JY, Takahashi DY, Ghazanfar AA. Cooperative vocal control in marmoset monkeys via vocal feedback. Journal of neurophysiology. 2015;114(1):274–83. doi: 10.1152/jn.00228.2015. PubMed PMID: WOS:000358010600025.

62. Jürgens U. The Neural Control of Vocalization in Mammals: A Review. Journal of Voice. 2009;23(1):1–10. doi: 10.1016/j.jvoice.2007.07.005.

63. Holstege G, Subramanian HH. Two different motor systems are needed to generate human speech. Journal of Comparative Neurology. 2016;524(8):1558–77. doi: 10.1002/cne.23898.

64. Hage SR, Nieder A. Dual Neural Network Model for the Evolution of Speech and Language. Trends in neurosciences. 2016;39(12):813–29. doi: 10.1016/j.tins.2016.10.006.

65. Jurgens U. Neural pathways underlying vocal control. Neuroscience and biobehavioral reviews. 2002;26(2):235–58. PubMed PMID: 11856561.

66. Alexander L, Wood CM, Gaskin PLR, Sawiak SJ, Fryer TD, Hong YT, et al. Over-activation of primate subgenual cingulate cortex enhances the cardiovascular, behavioral and neural responses to threat. Nature communications. 2020;11(1). doi: 10.1038/s41467-020-19167-0.

67. Zeredo JL, Quah SKL, Wallis CU, Alexander L, Cockcroft GJ, Santangelo AM, et al. Glutamate Within the Marmoset Anterior Hippocampus Interacts with Area 25 to Regulate the Behavioral and Cardiovascular Correlates of High-Trait Anxiety. The Journal of Neuroscience. 2019;39(16):3094–107. doi: 10.1523/jneurosci.2451-18.2018.

68. Jürgens U. The role of the periaqueductal grey in vocal behaviour. Behavioural brain research. 1994;62(2):107–17. doi: 10.1016/0166-4328(94)90017-5.

69. Tschida K, Michael V, Takatoh J, Han B-X, Zhao S, Sakurai K, et al. A Specialized Neural Circuit Gates Social Vocalizations in the Mouse. Neuron. 2019;103(3):459–72.e4. doi: 10.1016/j.neuron.2019.05.025.

70. Lu CL, Jurgens U. Effects of chemical stimulation in the periaqueductal gray on vocalization in the squirrel monkey. Brain Res Bull. 1993;32(2):143–51. PubMed PMID: 8102315.

71. Mateo C, Knutsen PM, Tsai PS, Shih AY, Kleinfeld D. Entrainment of Arteriole Vasomotor Fluctuations by Neural Activity Is a Basis of Blood-Oxygenation-Level-Dependent “Resting-State” Connectivity. Neuron. 2017;96(4):936–48.e3. doi: 10.1016/j.neuron.2017.10.012.

72. Ryan MJ. Energy, Calling, and Selection. American Zoologist. 1988;28(3):885–98. doi: 10.1093/icb/28.3.885.

73. Prestwich KN. The Energetics of Acoustic Signaling in Anurans and Insects. American Zoologist. 1994;34(6):625–43. doi: 10.1093/icb/34.6.625.

74. Gillooly JF, Ophir AG. The energetic basis of acoustic communication. Proceedings Biological sciences / The Royal Society. 2010;277(1686):1325–31. doi: 10.1098/rspb.2009.2134. PubMed PMID: 20053641; PubMed Central PMCID: PMC2871947.

75. Thomas R, Goldsmith A, Cuthill I, Lidgate H, Burdett Proctor S, Cosgrove D. The trade-off between singing and mass gain in a daytime-singing bird, the European robin. Behaviour. 2003;140(3):387–404. doi: 10.1163/156853903321826693.

76. Berg ML, Beintema NH, Welbergen JA, Komdeur J. Singing as a handicap: the effects of food availability and weather on song output in the Australian reed warblerAcrocephalus australis. Journal of Avian Biology. 2005;36(2):102–9. doi: 10.1111/j.0908-8857.2005.03285.x.

77. Voigt C, Leitner S, Gahr M. Seasonal Changes in the Song Pattern of the Non-Domesticated Island Canary (Serinus Canaria) a Field Study. Behaviour. 2001;138(7):885–904. doi: 10.1163/156853901753172700.

78. Garamszegi LZ, Moreno J, Moller AP. Avian song complexity is associated with high field metabolic rate. Evolutionary Ecology Research. 2006;8(1):75–90.

79. Bachman GC, Chappell MA. The energetic cost of begging behaviour in nestling house wrens. Animal Behaviour. 1998;55(6):1607–18. doi: 10.1006/anbe.1997.0719.

80. Wang TA, Yu YV, Govindaiah G, Ye X, Artinian L, Coleman TP, et al. Circadian Rhythm of Redox State Regulates Excitability in Suprachiasmatic Nucleus Neurons. Science. 2012;337(6096):839–42. doi: 10.1126/science.1222826.

81. Niven JE, Laughlin SB. Energy limitation as a selective pressure on the evolution of sensory systems. Journal of Experimental Biology. 2008;211(11):1792–804. doi: 10.1242/jeb.017574.

82. Mergenthaler P, Lindauer U, Dienel GA, Meisel A. Sugar for the brain: the role of glucose in physiological and pathological brain function. Trends in neurosciences. 2013;36(10):587–97. doi: 10.1016/j.tins.2013.07.001.

83. Li-Byarlay H, Rittschof CC, Massey JH, Pittendrigh BR, Robinson GE. Socially responsive effects of brain oxidative metabolism on aggression. Proceedings of the National Academy of Sciences. 2014;111(34):12533–7. doi: 10.1073/pnas.1412306111.

84. Bass AH, Clark CW. The Physical Acoustics of Underwater Sound Communication. 2003;16:15–64. doi: 10.1007/0-387-22762-8_2.

85. Timmer J. Modeling Noisy Time Series: Physiological Tremor. International Journal of Bifurcation and Chaos. 2011;08(07):1505–16. doi: 10.1142/s0218127498001157.

86. Friston KJ, Benayoun M, Cowan JD, van Drongelen W, Wallace E. Avalanches in a Stochastic Model of Spiking Neurons. Plos Comput Biol. 2010;6(7):e1000846. doi: 10.1371/journal.pcbi.1000846.

87. Dornic I, Chaté H, Muñoz MA. Integration of Langevin Equations with Multiplicative Noise and the Viability of Field Theories for Absorbing Phase Transitions. Physical review letters. 2005;94(10). doi: 10.1103/PhysRevLett.94.100601.

88. Danforth CM, Yorke JA. Making Forecasts for Chaotic Physical Processes. Physical review letters. 2006;96(14). doi: 10.1103/PhysRevLett.96.144102.

89. Strogatz SH. Nonlinear dynamics and chaos: with applications to physics, biology, chemistry, and engineering: Westview press; 2014.

